# Development of an amplicon-based high throughput sequencing method for genotypic characterisation of norovirus in oysters

**DOI:** 10.1101/2022.12.23.521849

**Authors:** Amy H Fitzpatrick, Agnieszka Rupnik, Helen O’Shea, Fiona Crispie, Paul D. Cotter, Sinéad Keaveney

## Abstract

Norovirus is a highly diverse RNA virus often implicated in food-borne outbreaks, particularly shellfish. Shellfish are filter feeders, and when harvested in bays exposed to wastewater overflow or storm overflows, they can harbour various pathogens, including human pathogenic viruses. The application of Sanger or amplicon-based High Throughput Sequencing (HTS) technologies to identify human pathogens in shellfish faces two main challenges i) distinguishing multiple genotypes/variants in a single sample and ii) low concentrations of norovirus RNA. Here we have assessed the performance of a novel norovirus capsid amplicon HTS method. We generated a panel of spiked oysters containing various norovirus concentrations with different genotypic compositions. Several DNA polymerase and Reverse Transcriptase (RT) enzymes were compared, and performance was evaluated based on i) the number of reads passing quality filters per sample, ii) the number of correct genotypes identified, and iii) the sequence identity of outputs compared to Sanger-derived sequences. A combination of the reverse transcriptase LunaScript and the DNA polymerase AmpliTaq Gold provided the best results. The method was then employed, and compared with Sanger sequencing, to characterise norovirus populations in naturally contaminated oysters.

**Importance:** While foodborne outbreaks account for approximately 14% of norovirus cases (Verhoef L, Hewitt J, Barclay L, Ahmed S, Lake R, Hall AJ, Lopman B, Kroneman A, Vennema H, Vinjé J, Koopmans M. 2015. 1999-2012. Emerg Infect Dis 21:592–599), we do not have standardised high-throughput sequencing methods for genotypic characterisation in foodstuffs. Here we present an optimised amplicon high-throughput sequencing method for the genotypic characterisation of norovirus in oysters. This method can accurately detect and characterise norovirus at concentrations typically detected in oysters. It will permit the investigation of norovirus genetic diversity in complex matrices and contribute to ongoing surveillance of norovirus in the environment.

## Introduction

In norovirus outbreaks associated with the consumption of shellfish, clinical and shellfish samples collected during the outbreak investigation are often subjected to nucleic acid Sanger sequencing for source attribution. The application of Sanger sequencing in this scenario is cumbersome in shellfish due to the need for the cloning of PCR isolates to capture the genetic diversity in a single sample (1–3) and generally only allows for low throughput analysis. High-throughput sequencing (HTS) permits low-cost sequencing, outputs high-throughput data and captures considerable nucleotide diversity, as demonstrated with 16S rRNA amplicon-based HTS analysis of bacteriomes. In contrast to Sanger sequencing, HTS technologies can also resolve multiple sequences per amplicon without cloning isolates in environmental samples.

A few studies have applied HTS-based methods for the detection and characterisation of norovirus in complex matrices such as food or wastewater (4–16). Notwithstanding these studies, the performance of the various methods, whether it be shotgun metagenomics, capture probe hybridisation, long-read sequencing or amplicon-based HTS, has varied. While shotgun metagenomics permits a less biased approach to sequencing, sequencing depth is often insufficient to characterise the norovirus present due to the presence of nucleic acid from other sources, even after rRNA removal or polyA tail enrichment (7, 8, 11, 12, 14). Capture probe hybridisation can enrich viral sequences, which could be helpful in challenging matrices. However, current market options are expensive and do not necessarily enrich the regions used for dual-genotyping norovirus, as panels are designed to broadly target viruses rather than specific viral families (7–9, 17). Long-read sequencing methods, such as those provided by PacBio and Oxford Nanopore Technologies (ONT), also face challenges in obtaining sufficient sequencing depth in complex matrices. Indeed, their outputs are typically lower than those from short-read platforms. ONT combined with adaptive sampling has had limited success in food samples (7), while, to date, long-read ONT sequencing of norovirus amplicons has not been successful (17). Despite these challenges, several recent studies have demonstrated the capability of various amplicon HTS assays for the genotypic characterisation of norovirus in shellfish (5, 17), with similar success observed in berry samples (6). However, these studies focused on application rather than optimisation and did not confirm HTS results with the gold standard Sanger methods. Due to the high degree of underreporting of norovirus cases, particularly non-nosocomial cases in healthy populations (18, 19), samples tend to come from food-borne outbreaks requiring source attribution or chronic nosocomial cases. This skews our understanding of norovirus genotypes circulating in local populations and will limit effective vaccine production in the future and management of clinical cases (20). Considering the necessity of capturing the norovirus sequences that permit genotypic characterisation in complex samples with low concentrations of viral RNA, this study has focused on optimising amplicon-based HTS methods.

Despite its potential value, amplicon HTS can introduce bias at multiple steps in the process, most notably RNA extraction and amplification of the target DNA. Bias during PCR cycling is impacted by choice of primers in that they are designed to target a conserved area of a chosen genome (21, 22). Currently, norovirus taxonomic assignment relies on dual genotyping based on the RNA- dependent RNA polymerase RdRp and VP1 gene (23). However, historically genotypic characterisation has focused on the VP1, also known as the capsid (24, 25). As a single-stranded RNA (ssRNA) virus, norovirus has a high mutation rate, estimated to be 5.40x10^-3^–2.23x10^-4^ nt substitutions/site/year for the VP1 encoding region (26). For example, SARS-CoV-2 has estimated 8.066x10^-4^ nt substitutions/site/year for the S gene (27). Moreover, noroviruses are genetically diverse, with a relatively low shared nucleotide identity of approximately 63% across commonly sequenced regions such as region C and the breakpoint of the RdRp-VP1 (23). Thus, designing suitable primers to capture norovirus’s existing and potential diversity is challenging. Various primers have been used for molecular characterisation, with most reference laboratories targeting the most conserved region of the genome (ORF1-ORF2 junction) and using degenerate primers that can tolerate sequence mismatches (25, 28). The regions typically targeted for molecular characterisation result in amplicons ranging from 113 bp to 587 bp in length (29). This study generated amplicons using primers targeting Region C (Figure 5), which yielded a 340 bp amplicon suitable for 300 bp paired-end sequencing. These primers are highly degenerate and can capture a broad range of norovirus genotypes.

The objective of this study was to establish a capsid amplicon-based HTS method for norovirus genotyping in shellfish. Critical criteria for a successful assay were as follows: i) accurate characterisation of genotypes in samples with concentrations of norovirus RNA typically observed in outbreaks and ii) accurate characterisation of multiple genotypes, and iii) superior performance to Sanger sequencing for application in surveillance-based studies. Reverse transcription (RT) and DNA polymerase enzymes are known to impact HTS outputs from quality to classification accuracy (30–33), therefore, selected enzyme combinations were compared in spiked and naturally contaminated samples.

The optimised amplicon-based HTS method and traditional Sanger sequencing method were successfully applied to a panel of naturally contaminated oysters, with a strong agreement between conventional and novel techniques.

## Results

The method was optimised with clinical samples and oysters either spiked with clinical material or naturally contaminated samples. By sequencing previously characterised clinical and spiked samples, method performance could be assessed in terms of its sensitivity and specificity.

### Characterisation of clinical positive control samples

To facilitate the subsequent assessment of the accuracy of HTS-based approaches, it was necessary first to source clinical samples containing specific norovirus genotypes. RT-qPCR and Sanger sequencing were used to characterise these clinical samples (Table 1), which were positive for one genotype per sample and contained a high concentration of norovirus RNA. These samples were subsequently used to spike oyster samples.

**Table 1:** Sanger sequencing results for clinical samples used in the proof-of-concept library and spiking experiments. Norovirus (NoV) detection is reported in genome copies per microlitre (gc/µL) for GI and GII. Genotypic characterisation of Sanger sequences from each sample was performed using NoroNet.

| Sample ID | NoV GI gc/μL | NoV GII gc/μL | Genotype NoroNet |
| --- | --- | --- | --- |
| FM011 |  | 1.03<sup>04</sup> | GII.3 |
| FM012 |  | 1.67<sup>03</sup> | GII.2 |
| FM018 |  | 1.18<sup>05</sup> | GII.4 Sydney 2012 |
| FM022 | 5.47<sup>03</sup> |  | GI.4 |
| FM023 |  | 3.23<sup>06</sup> | GII.4 |
| FM026 | 3.26<sup>02</sup> |  | GI.9 |

### Impact of enzyme combinations on genotype detection by HTS

Three reverse transcriptase and DNA polymerase enzyme combinations were applied to clinical and spiked shellfish samples. Table 2 provides the genotypes detected in the various sequencing experiments (1–3) by the enzyme combinations. Samples in experiment one included the clinical samples (Table 1) and spiked shellfish samples that were prepared using material from the clinical samples at various concentrations and combinations (Table 9). An additional spiked shellfish sample was sourced for a comprehensive RT and DNA polymerase comparison in experiment 2. The optimised method was applied to a panel of naturally contaminated samples in experiment 3, using LunaScript with AmpliTaq Gold.

**Table 2:** Genotypes detected using amplicon HTS across all sample types prepared with the selected RT and DNA polymerase enzymes. Genotypic characterisation of HTS Operational Taxonomic Units (OTUs) was performed using NoroNet.

| Enzyme combination | Genotype NoroNet |
| --- | --- |
| SuperScript II / AmpliTaq Gold | GI.3, GI.4, GI.7, GI.9, GII.2, GII.3, GII.4, GII.4 New Orleans 2009, GII.4 Sydney 2012, GII.6 |
| SuperScript II / Kapa HiFi | GI.3, GI.4, GI.7, GI.9, GII.2, GII.3, GII.4, GII.4 New Orleans 2009, GII.4 Sydney 2012, GII.6 |
| SuperScript II / Kapa Robust | GI.3, GI.4, GI.7, GI.9, GII.2, GII.3, GII.4, GII.4 New Orleans 2009, GII.4 Sydney 2012, GII.6 |
| SuperScript IV / AmpliTaq Gold | GI.3, GI.7, GII.6, GII.7 |
| SuperScript IV / Kapa HiFi | GI.3, GI.7, GII.6, GII.7 |
| SuperScript IV / Kapa Robust | GI.1, GI.3, GI.7, GII.2, GII.6, GII.7 |
| LunaScript / AmpliTaq Gold | GI.1, GI.3, GI.7, GI.9, GII.13, GII.14, GII.4 Sydney 2012, GII.6, GII.7 |
| LunaScript / Kapa HiFi | GI.1, GI.3, GI.7, GII.6 |
| LunaScript / Kapa Robust | GI.3, GI.7, GII.4 Sydney 2012, GII.6 |

**Table 9:**
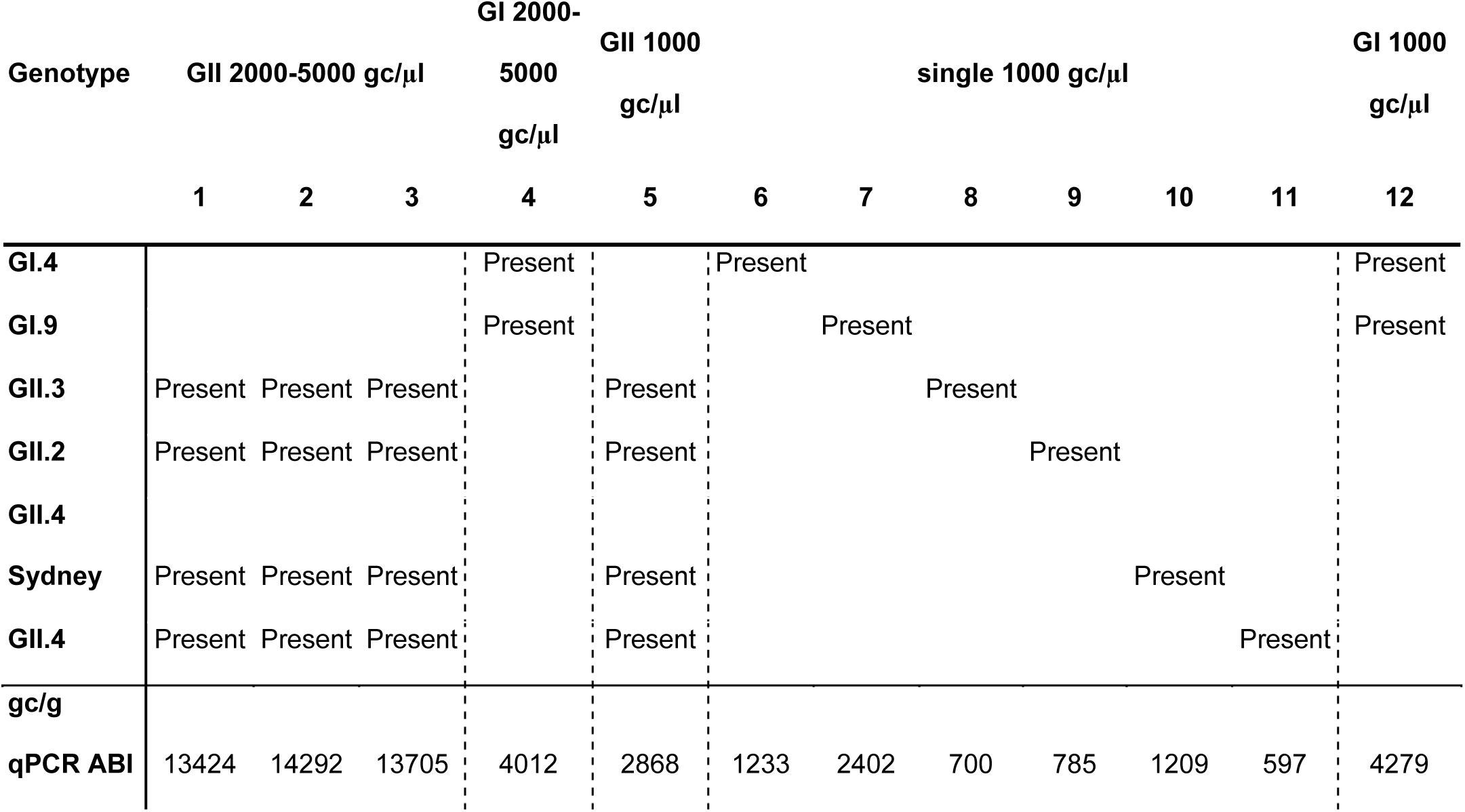
Spiked samples using in experiment one

### Enzyme combination impacts the quality of high throughput sequencing reads

Selected reverse transcriptase enzymes (RT) and DNA polymerases were compared to understand if method optimisation could improve the quality of sequences obtained. Moloney murine leukaemia virus (MMLV) derived RT SuperScript II, and SuperScript IV were evaluated alongside an *in-silico* designed RT LunaScript. The polymerases were selected because AmpliTaq Gold is widely used for RNA virus sequencing; Kapa HiFi has a low error rate and is recommended for use within 16S amplicon HTS protocols on Illumina platforms. In contrast, KAPA2G Robust has a greater tolerance for PCR inhibitors and is recommended for use with challenging samples.

Results were compared based on Phred quality scores, used to indicate the measure of base quality in DNA sequencing and expected errors, which are the sum of error probabilities over the length of the read. As per Figure 1. A, mean Phred scores obtained (spiked shellfish) were significantly different when assessed using a Kruskal-Wallis test (p-value 2.96^-12^, chi-squared = 425.56). RT and DNA polymerases are arranged from highest to lowest median Phred score, with LunaScript AmpliTaq Gold generating sequences with the highest Phred scores and LunaScript Kapa HiFi generating sequences with the lowest Phred scores. In Figure 1. B, expected errors (EE) were significantly different using a Kruskal-Wallis (p-value < 2.2^-16^, chi-squared = 1971.1). LunaScript AmpliTaq Gold generated sequences with the lowest mean EE, while Luna Kapa Robust generated sequences with the highest mean EE.

**Figure 1.**
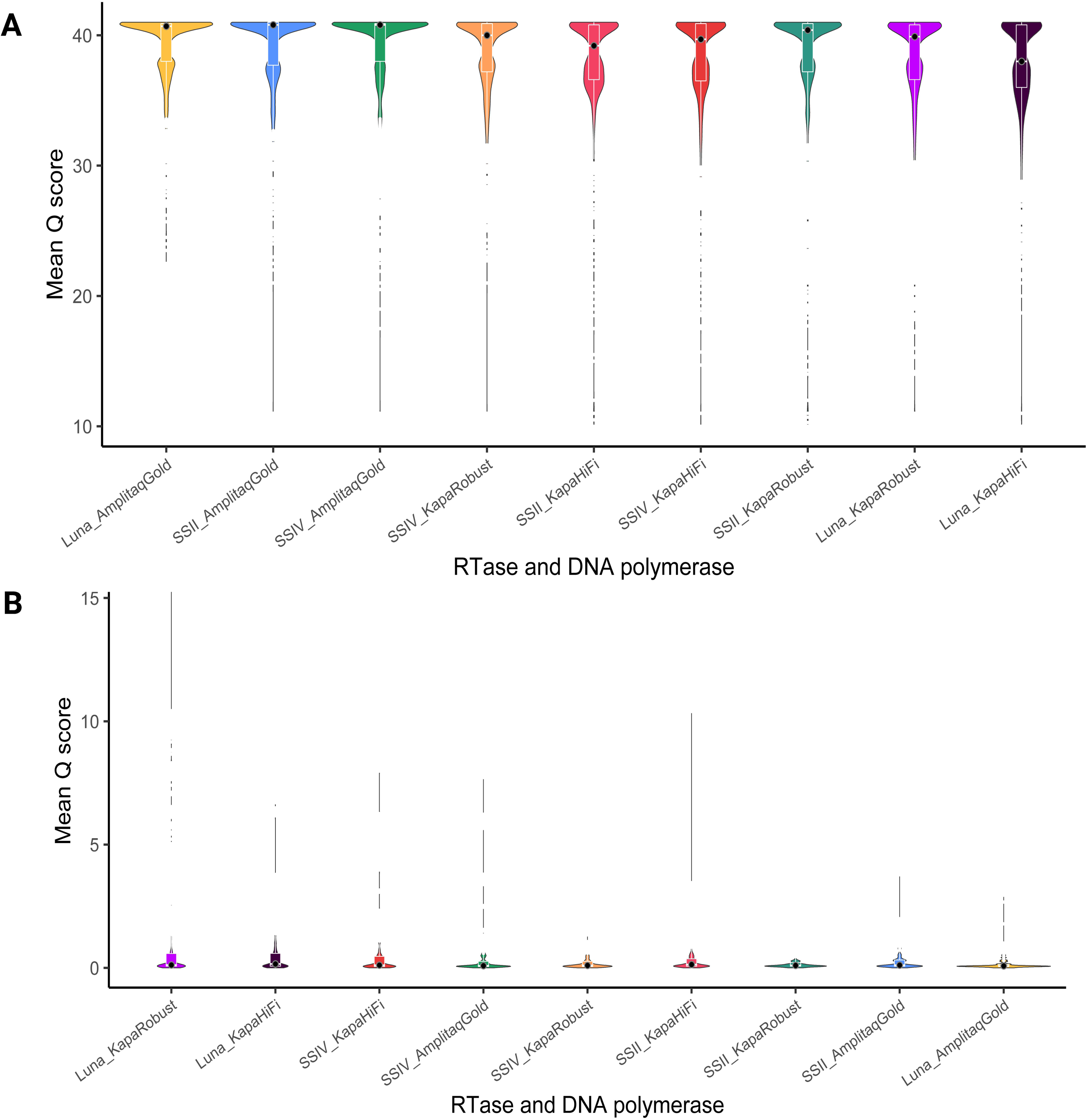
Violin plots with (A) an internal boxplot of the mean Phred score (Q) obtained in spiked shellfish and naturally contaminated oysters. RT and DNA polymerase combinations are ordered from highest mean Phred score to lowest (left to right). (B) Violin plots with internal boxplot of the mean expected errors (EE). RT and DNA polymerase combinations are ordered from the highest mean expected error score to the lowest (left to right).

A Factor Analysis of Mixed Data (FAMD) was performed to determine if RT or DNA polymerase enzyme contributed to the differences in performance in terms of quality (Figure 1). A FAMD works as a principal components analysis (PCA) for quantitative variables and a multiple correspondence analysis (MCA) for qualitative variables, allowing us to understand the relationship between numeric outcomes such as Phred score and factors such as DNA polymerase enzymes. In Figure 2. A, DNA polymerase enzymes contributed to 53% and 54% of the variation observed in dimensions 1 and 2, respectively; however, the overall cos2 value was low <0.3. A low cos2 indicates that the principal component does not perfectly represent DNA polymerase enzymes, i.e., other factors contribute to the variance observed. RT enzymes explained 22% and 49% of the variation observed in dimensions 1 and 2, respectively, though the cos2 value was low <0.3.

**Figure 2.**
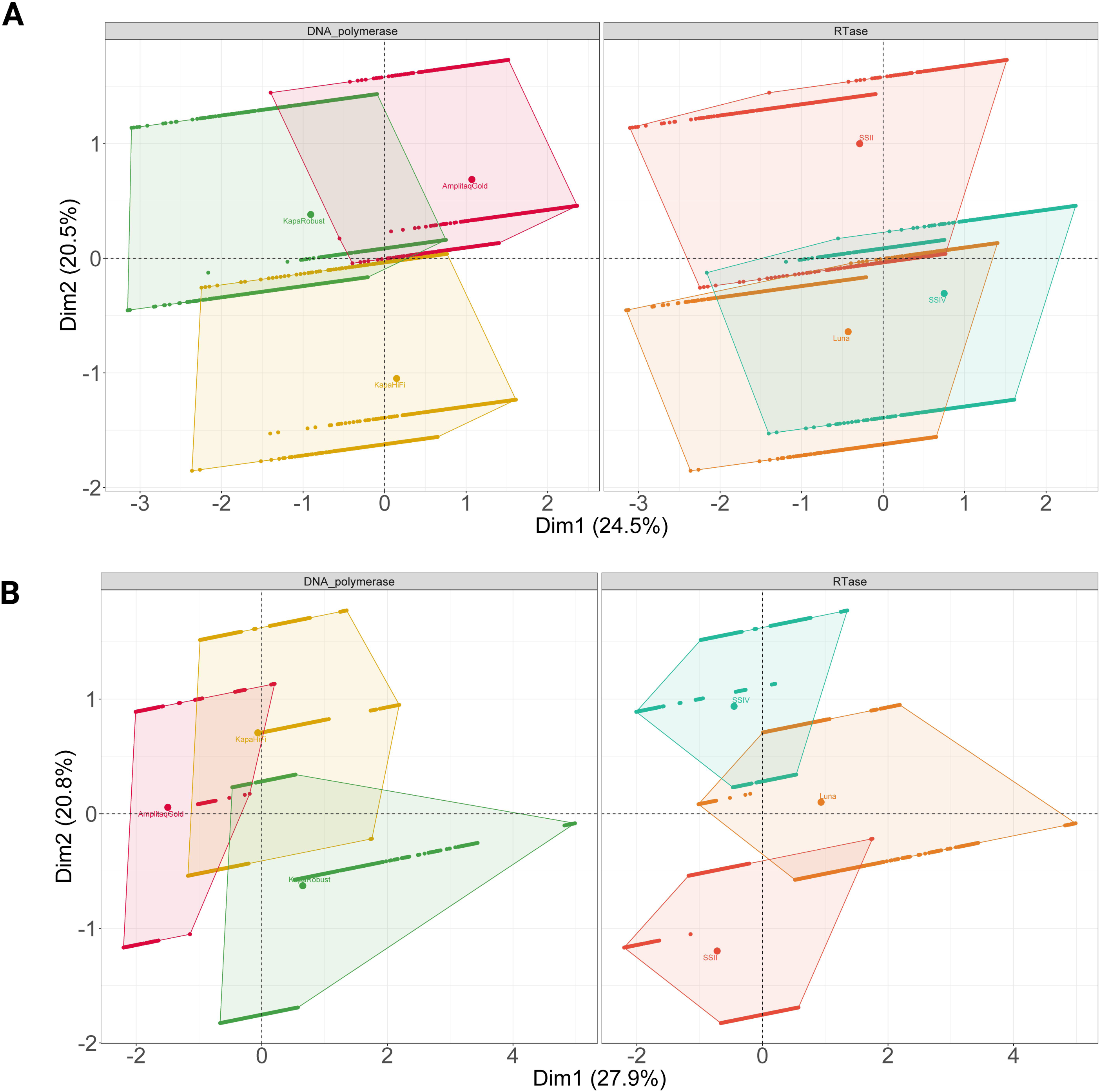
A Factor analysis with mixed data (FAMD) biplot demonstrates the variance-maximising distribution patterns of the total Mean Phred scores in the map space and their clustering patterns based on DNA polymerase and RT enzyme. B. FAMD biplot for DNA polymerase demonstrates the variance-maximising distribution patterns of the total Mean Expected Errors in the map space and their clustering patterns based on DNA Polymerase and RT enzyme.

In summary, DNA polymerase contributed to observed differences in Mean Phred score to a greater extent than the RT enzyme. On the other hand, as presented in Figure 2. B, DNA polymerase returned 42% and 35%, while RT enzyme returned 39% and 68% for dimensions 1 and 2, respectively. Cos2 values for both reagents were low, <0.3. Accordingly, mean EE was influenced by RT more than DNA polymerase enzymes. Therefore, RT enzyme choice has a greater impact on expected mean errors, while DNA polymerase has a greater effect on mean Phred scores.

### DNA polymerase impacts the relationship between input genomic material, and the number of HTS reads obtained

A Kendall rank correlation coefficient was used to compare the agreement between the concentration of DNA following library preparation or the gc/g of norovirus as determined by RT-qPCR before the semi-nested PCR to the HTS reads passing quality control. There was a perfect agreement (>0.8) between the concentration of DNA following library preparation as quantified by fluorometric measures and the resulting reads passing quality control for SuperScript II AmpliTaq Gold shellfish samples spiked with a single norovirus genotype (Table 3) (34). These correlations were not statistically significant, likely owing to the small sample size. Oyster samples that were spiked with multiple genotypes and prepared with SuperScript II AmpliTaq Gold provided a perfect agreement (>0.8) between the concentration of DNA following library preparation as quantified by fluorometric measures and the resulting reads passing quality control at statistically significant levels (Table 3). Overall, there was a weaker agreement between gc/g of norovirus as determined by RT-qPCR with reads passing quality control.

**Table 3.**
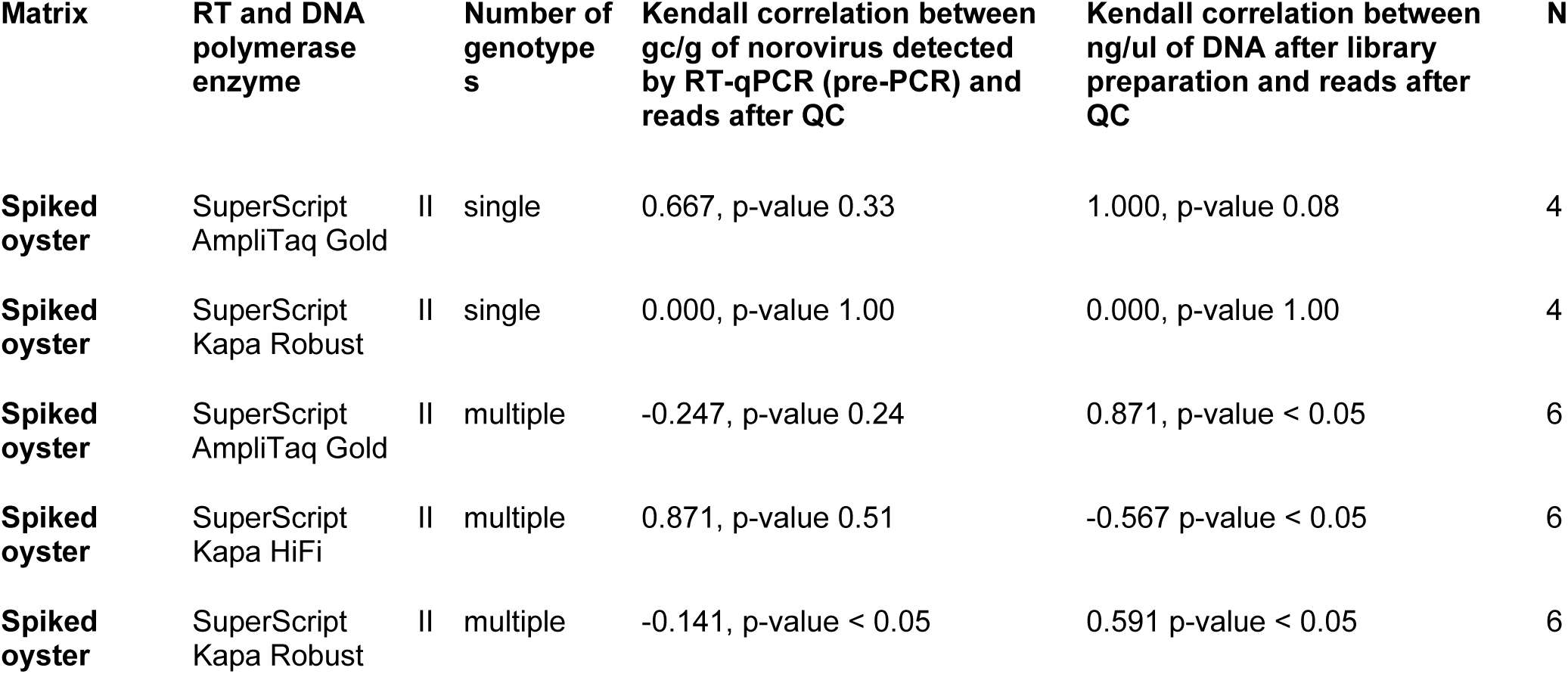
Kendall correlation between gc/g of norovirus detected by RT-qPCR prior to semi-nested PCR or ng/ul of DNA following semi-nested PCR and library preparation and reads after QC with HTS reads following quality control in spiked shellfish samples

Ultimately, in naturally contaminated samples (n=9), there was a moderate non-significant agreement of 0.481 (p-value 0.10) and 0.389 (p-value 0.18), respectively, between the gc/g of norovirus as determined by RT-qPCR and concentration of DNA following library preparation as quantified by fluorometric measures and the resulting reads passing quality control.

### Enzyme combination and technical triplicates can improve classification accuracy

A confusion matrix was used to investigate further the differences in performance between reverse transcription and DNA polymerase enzymes. As can be seen in Table 4, all enzyme combinations performed well when used with spiked shellfish samples. When the mock communities were prepared individually, all enzyme combinations returned perfect scores, apart from instances where SuperScript II was applied in combination with Kapa HiFi. For this library, GII.3 was not detected in mock community 8, present at 700 gc/g (single genotype). Libraries, where a spiked sample was prepared in triplicate from semi-nested PCR with LunaScript or SuperScript IV, provided perfect f-scores, in contrast to samples prepared with SuperScript II that did not provide perfect f-scores. The spiked sample contained GI.3, GI.7 and GII.6. GI.7 was not detected in samples prepared with SuperScript II in combination with AmpliTaq Gold or Kapa Robust. When the PCR products for norovirus GI and GII were combined, a loss in sensitivity was observed, i.e., GI.3 or GI.7 were not detected (Table 5). Similar trends were observed using the Jaccard index; see Supplementary, Figure 1.

**Table 4.** Performance of RT and DNA polymerase enzymes to genotype level classification as per a confusion matrix

| RT and DNA polymerase | Matrix | Technical replicates | accuracy | sensitivity | specificity | precision | recall | f-measure |
| --- | --- | --- | --- | --- | --- | --- | --- | --- |
| LunaScript AmpliTaq Gold | Spiked oyster | Triplicates | 1.00 | 1.00 | 1.00 | 1.00 | 1.00 | 1.00 |
| LunaScript Kapa HiFi | Spiked oyster | Triplicates | 1.00 | 1.00 | 1.00 | 1.00 | 1.00 | 1.00 |
| LunaScript Kapa Robust | Spiked oyster | Triplicates | 1.00 | 1.00 | 1.00 | 1.00 | 1.00 | 1.00 |
| SuperScript II AmpliTaq Gold | Spiked oyster | Individual | 1.00 | 1.00 | 1.00 | 1.00 | 1.00 | 1.00 |
|  | Spiked oyster | Triplicate | 0.86 | 0.83 | 1.00 | 1.00 | 0.83 | 0.91 |
| SuperScript II Kapa HiFi | Spiked oyster | Individual | 0.96 | 0.95 | 1.00 | 1.00 | 0.95 | 0.98 |
|  | Spiked oyster | Triplicate | 1.00 | 1.00 | 1.00 | 1.00 | 1.00 | 1.00 |
| SuperScript II Kapa Robust | Spiked oyster | Individual | 1.00 | 1.00 | 1.00 | 1.00 | 1.00 | 1.00 |
|  | Spiked oyster | Triplicate | 0.86 | 0.83 | 1.00 | 1.00 | 0.83 | 0.91 |
| SuperScript IV AmpliTaq Gold | Spiked oyster | Triplicate | 1.00 | 1.00 | 1.00 | 1.00 | 1.00 | 1.00 |
| SuperScript IV Kapa HiFi | Spiked oyster | Triplicate | 1.00 | 1.00 | 1.00 | 1.00 | 1.00 | 1.00 |
| SuperScript IV Kapa Robust | Spiked oyster | Triplicate | 1.00 | 1.00 | 1.00 | 1.00 | 1.00 | 1.00 |

**Table 5.**
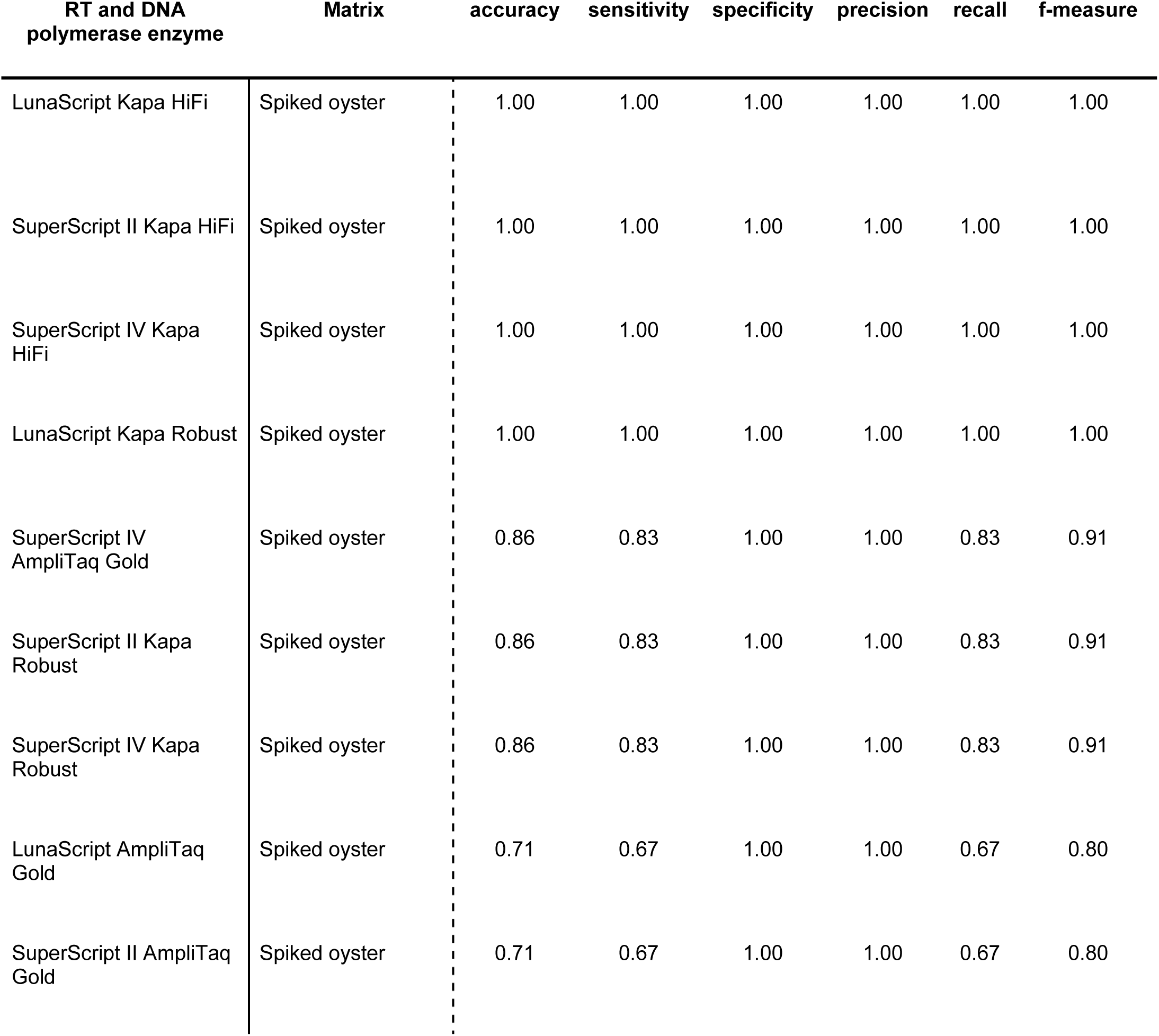
Performance of RT and DNA polymerase enzymes in pooled GI and GII amplicons to genotype level classification as per a confusion matrix

| RT and DNA<br>polymerase enzyme | Matrix | accuracy | sensitivity | specificity | precision | recall | f-measure |
| --- | --- | --- | --- | --- | --- | --- | --- |
| LunaScript Kapa HiFi | Spiked oyster | 1.00 | 1.00 | 1.00 | 1.00 | 1.00 | 1.00 |
| SuperScript II Kapa HiFi | Spiked oyster | 1.00 | 1.00 | 1.00 | 1.00 | 1.00 | 1.00 |
| SuperScript IV Kapa HiFi | Spiked oyster | 1.00 | 1.00 | 1.00 | 1.00 | 1.00 | 1.00 |
| LunaScript Kapa Robust | Spiked oyster | 1.00 | 1.00 | 1.00 | 1.00 | 1.00 | 1.00 |
| SuperScript IV AmpliTaq Gold | Spiked oyster | 0.86 | 0.83 | 1.00 | 1.00 | 0.83 | 0.91 |
| SuperScript II Kapa Robust | Spiked oyster | 0.86 | 0.83 | 1.00 | 1.00 | 0.83 | 0.91 |
| SuperScript IV Kapa Robust | Spiked oyster | 0.86 | 0.83 | 1.00 | 1.00 | 0.83 | 0.91 |
| LunaScript AmpliTaq Gold | Spiked oyster | 0.71 | 0.67 | 1.00 | 1.00 | 0.67 | 0.80 |
| SuperScript II AmpliTaq Gold | Spiked oyster | 0.71 | 0.67 | 1.00 | 1.00 | 0.67 | 0.80 |

### Phylogenetic distance between expected and observed sequences was affected by DNA polymerase enzyme

UniFrac is a distance matrix that measures the phylogenetic distance between sets of taxa in a phylogenetic tree. The distance is defined as the fraction of the branch length of the tree that leads to descendants from either one environment or the other, but not both. Unweighted UniFrac methods were used to compare the phylogenetic distance between sequences generated by SuperScript II RT and one of the following DNA polymerases; AmpliTaq Gold, Kapa Robust and Kapa HiFi. It demonstrated that DNA polymerase and RT enzyme explained some variations in sequencing results. DNA polymerase enzymes contributed to 35% of the variation observed in spiked shellfish. A Pairwise PERMANOVA was performed and returned significant p-values <0.05. Spiked samples prepared with SuperScript II, SuperScript IV or LunaScript RT enzymes and AmpliTaq Gold DNA polymerase returned p-values of 0.83-0.92, indicating a high similarity between obtained and expected sequences, see Supplementary Figure 2.

RT enzymes contributed to 37% of the variation observed in spiked shellfish. A post hoc test on the UniFrac distance matrix was performed using Pairwise PERMANOVA with 999 permutations and returned adjusted p-values of 0.84-0.93, respectively. This indicated high similarity between expected and obtained sequences when prepared with the same DNA polymerase, see Supplementary Figure 3.

Custom BLASTn databases were used to assess the ability of the various protocols to return a 99% match to the previously obtained Sanger sequences. SuperScript II AmpliTaq Gold and SuperScript II Kapa Robust returned a 99% BLASTn match with bit-scores >500 and e-values < 0.001 for all expected genotypes. SuperScript II Kapa HiFi I failed to produce a 99% match for the GI.9 genotype; see Supplementary Table 1.

Overall, LunaScript and AmpliTaq Gold provided the most accurate, high-quality results based on quality metrics, classification accuracy and phylogenetic distance.

### HTS amplicon sequencing of the capsid region permitted the detection of additional genotypes from naturally contaminated shellfish samples compared to the conventional Sanger sequencing

Based on the preceding results, a library with naturally contaminated oysters was prepared using LunaScript in combination with AmpliTaq Gold for RT and the semi-nested PCR. Three naturally contaminated samples with varying concentrations of norovirus GII (see Table 6) were lysed and extracted for amplicon-based HTS and Sanger sequencing.

**Table 6:**
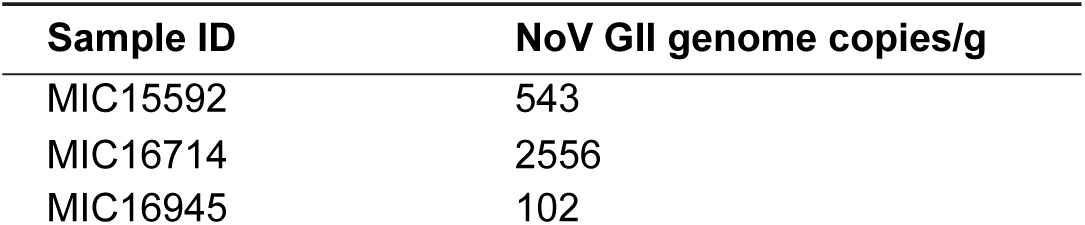
Naturally contaminated samples sequenced using Sanger and HTS

| Sample ID | NoV GII genome copies/g |
| --- | --- |
| MIC15592 | 543 |
| MIC16714 | 2556 |
| MIC16945 | 102 |

Sequences obtained using Sanger sequencing could not be genotyped using NoroNet. However, CaliciNet and the internal classifier provided strong concordance of genotype assignment with the MiSeq results, see Table 7. Technical triplicates introduced from the first round of RT-PCR provided strong agreement regarding the relative abundance observed for each genotype detected, see Supplementary Figure 4. More genotypes were detected using the amplicon HTS method than conventional Sanger sequencing of cloned variants. Three genotypes were detected using Sanger sequencing in MIC16714 (GII.6, GII.4 Sydney, GII.13), while an additional GII.7 was detected using amplicon HTS. In MIC15592, GII.14 was detected using Sanger sequencing, while amplicon HTS detected both GII.14 and GII.6.

**Table 7.** Norovirus genotypes detected using both Sanger sequencing and HTS amplicon sequencing

| Sample | Sanger with cloning of isolates | Amplicon HTS MiSeq |
| --- | --- | --- |
| MIC15592 | GII.14 | GII.14, GII.6 |
| MIC16714 | GII.6, GII.4 Sydney 2012, GII.13 | GII.4 Sydney 2012, GII.13, GII.6, GII.7 |
| MIC16945 | GII.6 | GII.6 |

## Discussion

In this study, we have demonstrated that Reverse Transcription and DNA polymerase enzymes impact HTS library quality. RT-qPCR data on norovirus genome copies/g was a moderate indicator of obtained HTS reads. The optimised extraction and semi-nested PCR method permitted the accurate detection of norovirus in naturally contaminated oysters when combined with a custom bioinformatic pipeline. This is an essential development for environmental virology. The application of HTS to genotype norovirus in contaminated foods has been constrained due to a lack of available methods.

This study focused on reverse transcription and semi-nested PCR steps for optimisation. In terms of the quality of the sequences observed, as measured by Phred score and expected errors, combined enzyme choice shaped score profiles (Figures 1 and 2). It has been well documented that the priming strategy and the RT enzyme can impact the reverse transcription (RT) of RNA to cDNA. However, previous norovirus HTS studies using custom hexamers reported no improvement in performance compared to random hexamers (8). While RT aims to produce cDNA that faithfully reflects the starting RNA sample, several studies indicate that the RT reaction can introduce large variability (35–37).

DNA polymerase contributed more to the variation observed for mean Phred scores, while RT enzymes contributed more to the variability observed for mean expected errors (Figure 2). In particular, Kapa HiFi and KAPA Robust had higher expected errors when combined with LunaScript (Figure 2. B); conversely, AmpliTaq Gold and LunaScript provided the lowest expected errors overall. This implies that RT and DNA polymerase combinations operate synergistically. Nonetheless, the literature on the mechanism behind varying RT and DNA polymerase enzyme performance is limited. The initial publication describing AmpliTaq Gold outlined its superior performance in complex sample types with low genomic input and/or multiple PCR assays (38). At the same time, KAPA Robust has been recommended for amplification in samples with high levels of inhibitors (39). AmpliTaq Gold has been widely applied in molecular virology (40–42), though there is limited literature evaluating its performance relative to other DNA polymerases. In this case, LunaScript combined with AmpliTaq Gold provided the highest quality sequences; however, previous studies in other complex sample types provide conflicting results (30, 32, 43–46). Several factors may contribute to the performance of RT and DNA polymerase enzymes, such as low input genomic material, RNA quality, matrix-specific factors, target-specific factors and DNA synthesis speed. PCR is a stochastic amplification process challenged by multiple templates, secondary structures and GC content (33, 47). The predicted hairpin structures in norovirus (48, 49) and the presence of multiple genotypes in shellfish challenge the development of any HTS applications. This warrants further study, as it is important to understand why performance variation is observed to optimise it.

Of note, AmpliTaq Gold provided high-quality sequences and optimal results in terms of classification accuracy when combined with specific RT enzymes, even though the semi-nested PCR assay with AmpliTaq Gold was performed with the highest number of total cycles. The number of PCR cycles is known to influence results. While a higher number of PCR cycles might increase the likelihood that rare molecules are observed, it can also skew abundance estimates by amplifying the biases (32, 50). However, this was not the case in this study. There are no comparison studies on LunaScript, as it has only recently been added to the market, but it is widely used for the RT step in the ARTIC SARS-COV-2 protocol (51). AmpliTaq Gold has a slower DNA synthesis rate than the other studied polymerases; Kapa HiFi and Kapa Robust. Furthermore, nanopore systems have demonstrated that slower translocation rates result in high accuracy (52); therefore, faster synthesis is not necessarily equally as accurate, especially in the case of highly diverse amplicons.

Various attempts have been made to optimise the workflow in terms of the wet-lab methodology developed. Notably, applying the ISO 15216:2017 method in combination with the optimised semi-nested PCR did not successfully amplify norovirus in naturally contaminated samples. Several modifications were required, including the concentration of the viral RNA by eluting it into a lower volume and increasing the input cDNA in the first round of the semi-nested PCR. This emphasises the importance of performing method development with the target matrix, i.e., in this case, naturally contaminated shellfish, as spiked shellfish samples performed well without modifications. In addition, the inclusion of technical triplicates incorporated in the various experiments resulted in improved results relative to instances where individual samples were used. This observation is consistent with previous work (17).

Furthermore, it was notable that the pooling of amplicons from norovirus GI and GII PCR assays before library preparation resulted in lower classification accuracy, with markedly fewer reads aligning to norovirus GI. While these steps increase the workload per biological sample, we find they are necessary for optimal HTS results. Enzyme choice (RT/DNA polymerase) did impact the accuracy of HTS of norovirus VP1 amplicons. Almost all combinations of enzymes returned perfect f-scores (1.00) when performed in triplicate apart from those treated with SuperScript II. Thus all genotypes known to be present in the samples were detected with no false positives. In the first experiment, clinical samples and spiked shellfish were sequenced; the presence of GI.9 was not detected using SuperScript II in combination with Kapa HiFi in a clinical sample. Indeed, the GI.9 sequence in question has four known nucleotide mismatches with the primers used in this study. No genotypes were missed using the LunaScript / AmpliTaq Gold enzyme combination in spiked samples. The high concordance between MiSeq amplicon HTS and conventional Sanger sequencing results supports method application in naturally contaminated shellfish.

An important consideration in choosing suitable samples to process using the outlined methodology is the detected norovirus gc/g as per ISO 15216:2017 (53). As per the moderate correlation between input gc/g and obtained HTS reads, we advise selecting samples greater than 100 gc/g for norovirus amplicon HTS (Table 6). Notwithstanding this recommendation, it has been observed in this study that some samples with a high concentration of viral RNA may fail to produce peaks of the expected size (2100 Bioanalyzer), while samples containing <300 gc/g may produce high-quality sequences. This is likely due to the quality and fragmentation of the norovirus RNA present in the shellfish at hand.

The semi-nested PCR targets the VP1 capsid region of norovirus, and the RT-qPCR targets a smaller overlapping region in the VP1. Amplification or sequencing of the full-length VP1 region has been used as a proxy for infectivity due to the hypothesis that an intact capsid infers an intact virus capable of initiating an infection (54–57). As observed in this study, spiked samples may behave differently from naturally contaminated oysters as the norovirus RNA is intact. In contrast, norovirus accumulated in shellfish could be degraded and fragmented by wastewater treatment processes (2, 58) and/or exposure to UV in the marine environment (59, 60). This supported the variation in the correlation between input material and obtained HTS reads from spiked to naturally contaminated samples (Table 3). Despite this observation, bioaccumulation experiments with fragmented norovirus RNA and clinical samples established that the intact virus was preferentially bioaccumulated over fragments of viral RNA and could survive up to two weeks (61). In a previous trial, norovirus remained infectious for up to 61 days in groundwater at room temperature. It persisted for up to 3 years based on RNase+ RT-qPCR assays (62), though recent publications utilising the Human Intestinal Enteroid (HIE) models have indicated a much shorter persistence of viable norovirus (63).

Furthermore, it has been well documented that noroviruses can be harboured within biofilms, resulting in increased persistence (64) and binding to histo blood group antigens (HBGA)-like molecules on enteric bacteria, increasing persistence and enhancing viral pathogenesis (65, 66). As the target regions for RT-qPCR of norovirus are < 100 bp, it is not surprising to observe less than a perfect agreement between the HTS reads for 340/344 bp amplicons from a semi-nested PCR and gc/g as per RT-qPCR amplicons. Therefore, it is challenging to define the probability that norovirus viral RNA detected by RT-qPCR in shellfish is an intact and/or infectious virus (67, 68).

While this study builds on previous work (5, 8, 67) and enhances the capacity for surveillance and outbreak response on a national level, there are limitations. Much of the work presented was performed with spiked samples, which are not necessarily representative of naturally contaminated shellfish due to the quality and concentration of the RNA. Unfortunately, RNA quality is difficult to assess in molluscs due to a hidden break in the 28S (69), making it challenging to obtain a RIN value. Therefore, we could not compare samples based on their RNA quality. Additionally, we could not represent the full genetic diversity of noroviruses due to limited access to clinical samples. Moreover, PCR-based sequencing will always be biased due to the choice of primers.

We hypothesise that updated primer sets would permit the detection of additional genotypes, particularly for norovirus GII.17, GII.3 and GI.3 (4, 21, 28, 70). While a confusion matrix was used to assess method performance, an inter-laboratory ring trial would provide a more realistic measure of method performance. Finally, clinical genotyping of norovirus relies on the RdRp and VP1 of norovirus (23, 71, 72). Ideally, the RdRp and VP1 should be amplified and sequenced in one amplicon, yet Illumina limitations concerning read length do not permit this. The application of Oxford Nanopore Technology or PacBio sequencing platforms would enable the sequencing of longer amplicons and merits investigation.

This study provides a fit-for-purpose protocol for the genotypic characterisation of norovirus in shellfish. We determined that Reverse Transcription and DNA polymerase enzyme choice, technical triplicates and an optimised RNA extraction procedure impact the quality and accuracy of HTS of norovirus amplicons. Wet-lab methodology optimisation is pivotal in moving the field from *ad-hoc* sequencing to accredited methods. The results provided here have wide-ranging implications for HTS study design. Establishing standardised and well-described HTS methods, from the wet lab to the bioinformatic analysis, is vital for building consensus in outbreak investigations across shared jurisdictions. The methods we present here can be applied for widespread surveillance of norovirus in complex samples such as shellfish or wastewater to expand our understanding of norovirus diversity.

## Methods

### Samples

To assess the performance metrics, a series of three different experiments were performed. An overview of the sequencing libraries is provided in Table 8.

**Table 8:** Overview of sequencing experiments

| Experiment | sample matrix | Technical replicates | No of biological samples | Total PCR products | Genotypes Expected |
| --- | --- | --- | --- | --- | --- |
| Proof of concept | clinical & spiked shellfish | Individual | 18 (6 clinical & 12 spiked shellfish samples) | 59 | GI.4, GI.9, GII.3, GII.2, GII.4 Sydney 2012, GII.4 |
| RT and DNA polymerase comparison | spiked & naturally contaminated shellfish | Triplicate | 2 | 98 | GI.3, GI.7, GII.6 |
| Application in naturally contaminated shellfish | naturally contaminated shellfish | Triplicate | 3 | 10 | unknown |

First, a proof-of-concept library was prepared and sequenced using SuperScript II and a selection of different DNA polymerases in both clinical (Table 1) and spiked samples (Table 9). A total of six stool samples were used as positive controls, while twelve matrix-specific mock communities were prepared.

Spiked and naturally contaminated samples were prepared in triplicate for a comprehensive RT and DNA polymerase enzyme comparison in the second experiment. The spiked sample was obtained from the proficiency testing scheme operated by the European Reference Laboratory for foodborne viruses (MIC200561). MIC200561 contained approximately 10000 copies of GI, and 1000 copies of GII (GI.3, GII.6, GII.7), while MIC180026 was a sample from a harvesting site collected in January 2018. It contained approximately 1000 gc/g GI and 4500 gc/g GII.

In the final experiment, three naturally contaminated oysters harvested in 2015 and 2016 were subject to an optimised protocol (RNA extraction to RT-PCR) to demonstrate the application of the method in the target samples, see Table 6.

### Preparation of Oyster and Faecal Samples for Norovirus Analysis

In line with ISO 15216-1:2017, oysters were tested for the presence of norovirus GI and GII (73). In brief, oysters were cleaned before shucking and dissecting 10 oysters per sample. The dissected digestive tissue (DT) was diced and combined was a sterile razor blade. Samples were lysed with 2 ml Proteinase K (100 g ml^-1^), followed by incubation and shaking at 37 °C for 60 minutes at 150 rpm. Samples underwent an additional incubation period of 15 minutes at 60 °C. Supernatants were retained for RNA extraction following centrifugation at 3000×g for 5 min.

For each stool sample, 0.5 ml PBS (Oxoid, UK) was added to a 2 ml tube containing between 2 g faecal material (neat) and vortexed vigorously. Then, 100 μl of the resuspended faecal material (neat) was transferred into a fresh tube containing 900 μl of PBS (10^-1^) and serial dilutions were prepared up to 10^-5^.

### Viral RNA extraction

NucliSENS® magnetic extraction reagents (bioMérieux) and the NucliSENS® EasyMag® extraction platform were used to extract RNA from 500 µl of DT supernatants. This was then eluted into 100 µl of elution buffer. RNA extracts were kept at − 80 °C until the RT-qPCR or semi-nested PCR analysis was carried out. A single negative extraction control (water) was performed.

### Determination of the Norovirus Concentration Using One-Step RT-qPCR

For samples where quantification of the norovirus RNA is provided, RT-qPCR was performed per ISO 15216-1:2017 (73). RT-qPCR analysis was performed using the Applied Biosystems AB7500 instrument (Applied Biosystems, Foster City, CA) and the RNA Ultrasense one-step RT-qPCR system (Invitrogen). The following were combined on a 96-well optical reaction plate to prepare the reaction mixture: 5 µl of RNA and 20 µl of the reaction mix containing 500 nM forward primer, 900 nM reverse primer, 250 nM sequence-specific probe, 1 × ROX reference dye and 1.25 µl of enzyme mix. Norovirus GI was detected using previously described primers QNIF4 (74), NV1LCR (75) and the TM9 probe (76), while QNIF2 (77), COG2R (78) and QNIFS probe (77) were used to detect norovirus GII. The internal process control mengovirus was detected using Mengo110, Mengo209 and Mengo147 probe (79). The plate was incubated at 55 °C for 60 min, 95 °C for 5 min, and then 45 cycles of PCR were performed, with 1 cycle consisting of 95 °C for 15 s, 60 °C for 1 min, and 65 °C for 1 min. All samples were analysed for norovirus GI and GII in duplicate. All control materials used in the RT-qPCR assays were prepared as previously described (80).

### Preparation of matrix-specific mock communities

A panel of norovirus matrix-specific mock communities were generated using the positive control material (Table 1). Based on the genome copies (gc) per µl of each genotype, the weight of the norovirus-positive faecal sample to be added to the negative digestive tissue for the desired ratio was calculated and spiked into the homogenised norovirus-negative oyster digestive tissue. The total norovirus concentration in each mock community ranged from 597 gc/g to 14292 gc/g of NoV GI or GII RNA, see Table 9.

### cDNA generation and semi-nested PCR

cDNA was generated using either SuperScript II, SuperScript IV or LunaScript RT as per the manufacturer’s protocols. Three DNA polymerases were evaluated for performance on the semi-nested PCR; AmpliTaq Gold, Kapa HiFi and Kapa Robust. For AmpliTaq Gold, the first round of nested PCR was prepared as follows, 5 µl cDNA was added to a 45 μl reaction mixture with a final concentration of 10 mM Tris- HCl (pH 8.3), 50 mM KCl, 20 μM of dNTPs, 2 μM of each primer (see Table 10), 2.5mM of MgCl2 and 2.5 U of AmpliTaq® DNA Polymerase (Applied Biosystems, USA). For the second round of PCR, the first round PCR product (5 µl) was added to 45 µl of a reaction mixture containing 10 mM Tris-HCl (pH 8.3), 50 mM KCl, 20 μM of dNTPs, 0.4 μM of each primer, 2.5 mM of MgCl2 and 2.5 U of AmpliTaq® DNA Polymerase.

**Table 10:**
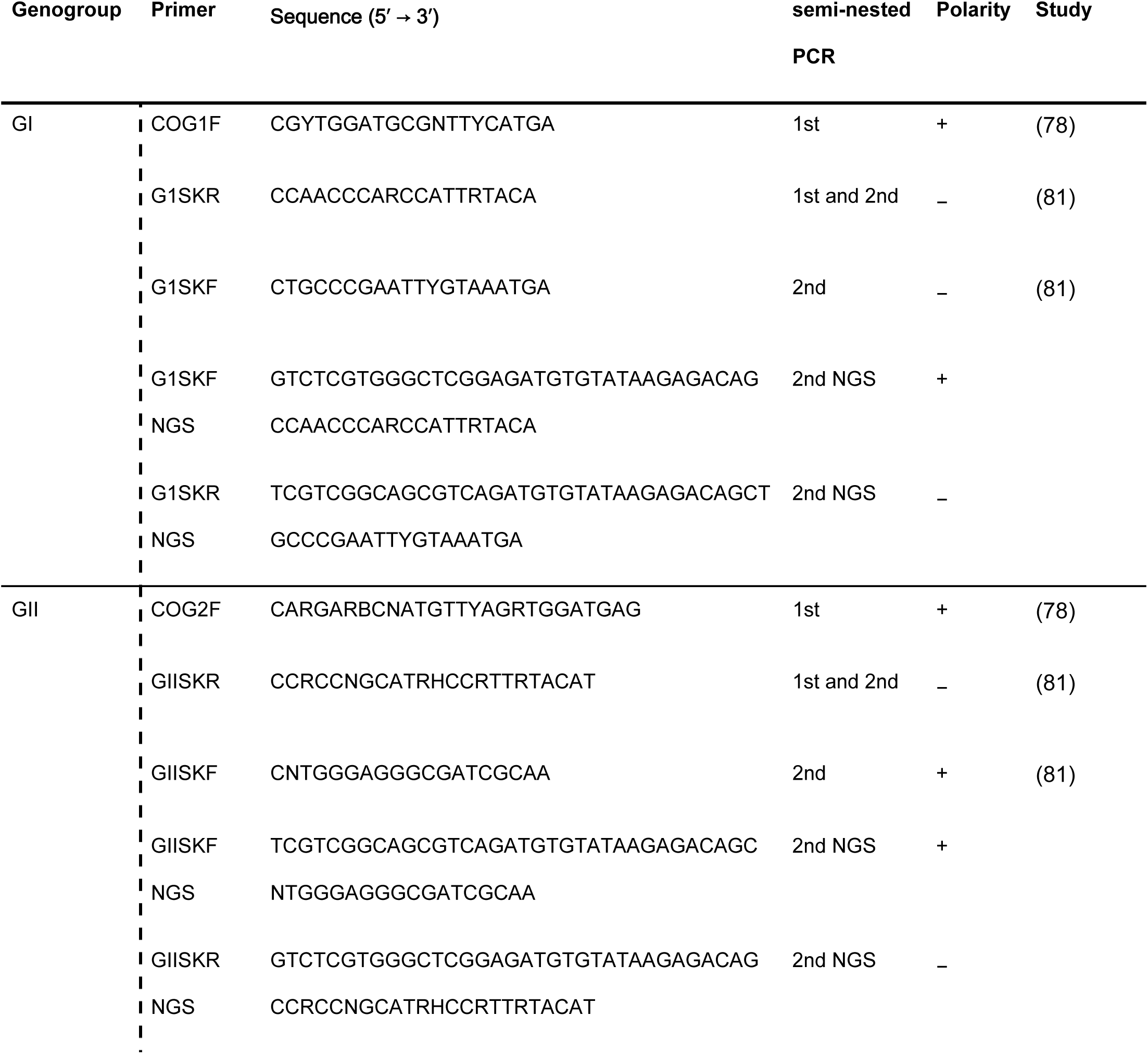
Primers used for amplification of VP1 capsid gene in Sanger and Illumina sequencing

For KAPA HiFi HotStart ReadyMix kit (Kapa Biosystems), the first round of PCR was prepared as follows: 5 µl cDNA with a final concentration of 10 μM of primers, 12.5 μl of KAPA HiFi HotStart ReadyMix and 2.5 μl of molecular grade biology water in a 25 μl reaction volume. For the second round of PCR, the first round PCR product (2.5 µl) was added to 22.5 µl of a reaction mixture containing a final concentration of 10 μM for primers, 12.5 μl of KAPA HiFi HotStart ReadyMix and 5 μl of molecular grade biology water. For KAPA2G Robust HotStart ReadyMix (Kapa Biosystems), the first round PCR was performed as follows: 5 µl cDNA with a final concentration of 10 μM of primers, 12.5 μl of KAPA2G Robust HotStart ReadyMix and 2.5 μl of molecular grade biology water. For the second round of PCR, the first round PCR product (1 µl) was added to 24 µl of a reaction mixture containing a final concentration of 10 μM for primers, 12.5 μl of KAPA2G Robust HotStart ReadyMix and 6.5 μl of molecular grade biology water. PCR conditions are described in Table 11.

**Table 11:**
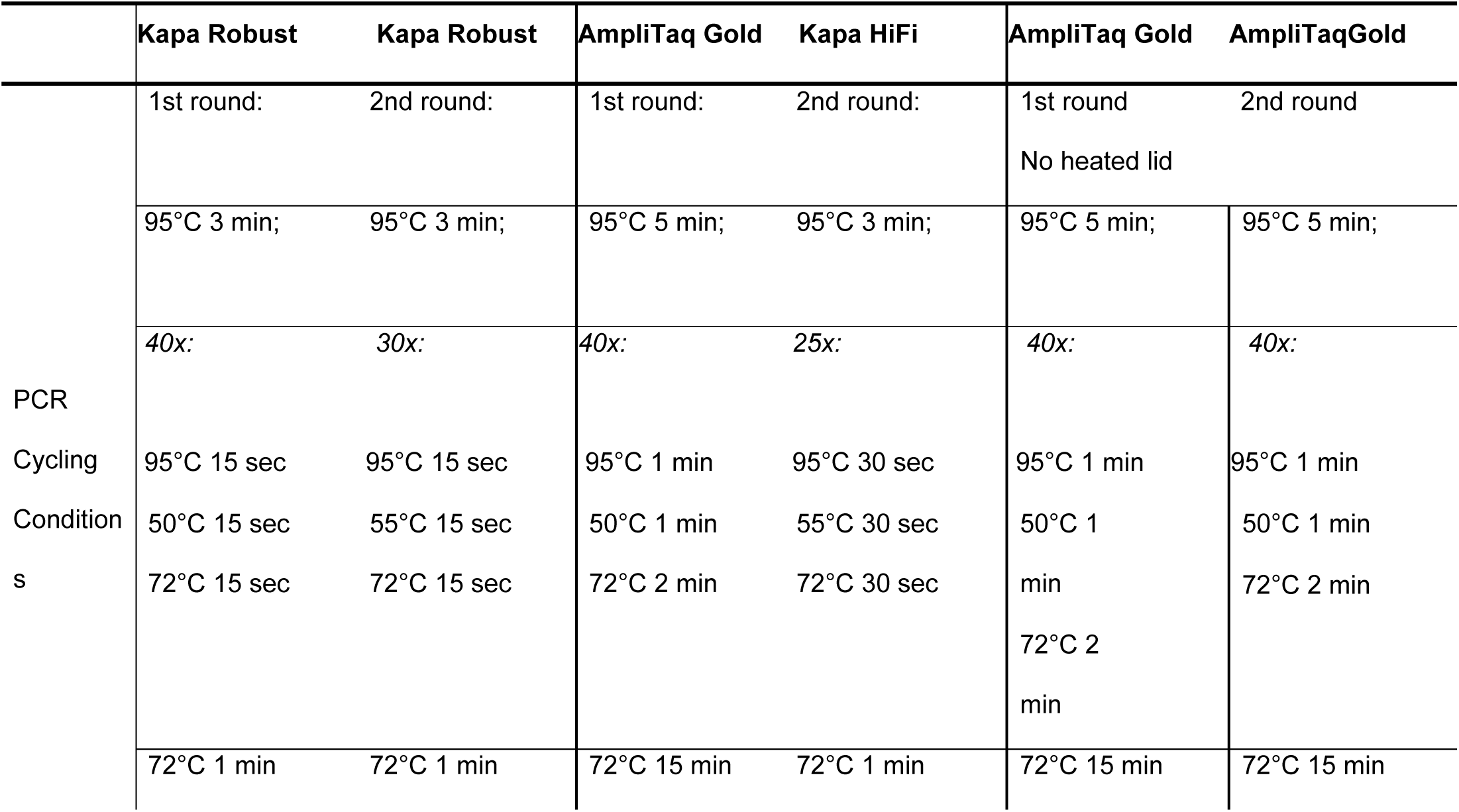
PCR conditions for the norovirus capsid region

|  | Kapa Robust | Kapa Robust | AmpliTaq Gold | Kapa HiFi | AmpliTaq Gold | AmpliTaqGold |
| --- | --- | --- | --- | --- | --- | --- |
|  | 1st round: | 2nd round: | 1st round: | 2nd round: | 1st round | 2nd round |
|  |  |  |  |  | No heated lid |  |
|  | 95°C 3 min; | 95°C 3 min; | 95°C 5 min; | 95°C 3 min; | 95°C 5 min; | 95°C 5 min; |
|  | 40x: | 30x: | 40x: | 25x: | 40x: | 40x: |
| PCR |  |  |  |  |  |  |
| Cycling | 95°C 15 sec | 95°C 15 sec | 95°C 1 min | 95°C 30 sec | 95°C 1 min | 95°C 1 min |
| Condition | 50°C 15 sec | 55°C 15 sec | 50°C 1 min | 55°C 30 sec | 50°C 1 | 50°C 1 min |
| s | 72°C 15 sec | 72°C 15 sec | 72°C 2 min | 72°C 30 sec | min | 72°C 2 min |
|  |  |  |  |  | 72°C 2<br>min |  |
|  | 72°C 1 min | 72°C 1 min | 72°C 15 min | 72°C 1 min | 72°C 15 min | 72°C 15 min |

PCR individual or triplicates were performed during the first round of semi-nested PCR, using priming sites as shown in Figure 3. The primers for Sanger sequencing and HTS (see NGS primers) are provided in Table 10. Second-round PCR products were visualised on a 1x TAE 2% agarose gel containing 5 μl of Ethidium Bromide for clinical and spiked shellfish samples. The Agilent High Sensitivity DNA Kit for naturally contaminated shellfish was used to visualise second-round PCR products for Bioanalyzer 2100 (Agilent Technologies).

**Figure 3:**
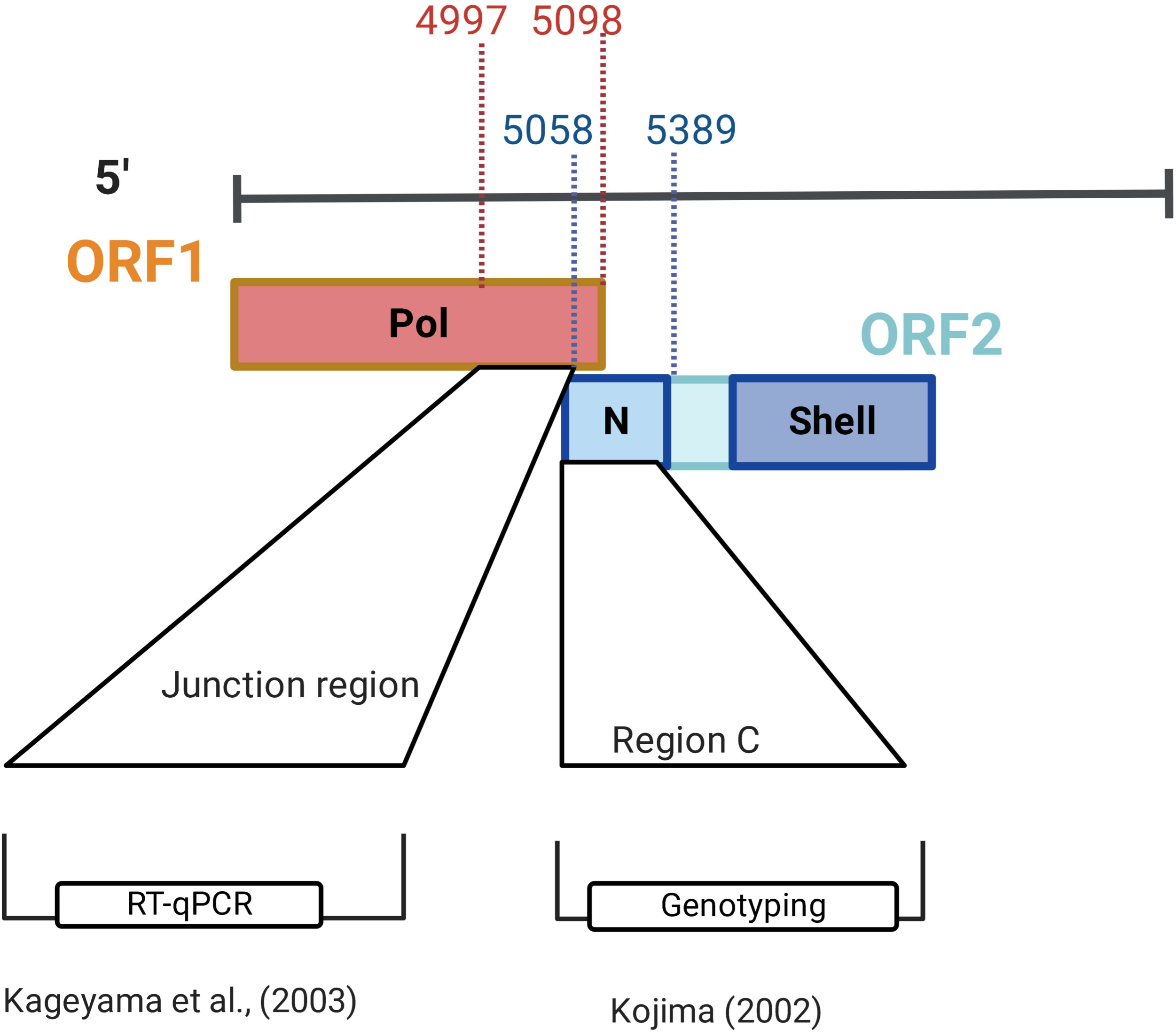
Regions of norovirus genome used for genotypic characterisation and detection by RT-qPCR.

### Cloning for Sanger sequencing

PCR amplicons were gel extracted using the QIAquick gel extraction kit and ligated into pGEM-T Easy plasmid (Promega). Vectors containing the PCR products were then cloned into chemically competent *E. coli*. Diluted and undiluted cells were plated on Luria Broth (LB) agar (Sigma, UK) containing XGal (20mg/ml), IPTG (100mM/ml) and Ampicillin (100mg/ml). Approximately ten to fifty colonies per sample were picked and purified using the QIAprep spin miniprep kit. PCR then confirmed the presence of the target DNA with M13 forward -20 (TGTAAAACGACGGCCAGT) and M13 reverse -27 primers (CAGGAAACAGCTATGAC) with Kapa Robust as per the manufacturer’s instructions. Sanger sequencing was used to obtain the nucleic acid sequences of cloned fragments.

### MiSeq library preparation and sequencing

Illumina sequencing adapters were incorporated into the second round PCR primers; see Table 10 for the sequences of the primers. PCR products were purified using Ampure XP beads (Beckmann Coulter) (0.8x bead:pool ratio) with elution of PCR products in 25 µl following the first bead clean-up. Cleaned-up PCR products were then indexed with the Nextera XT index kit (Illumina) following a modified 16S rRNA protocol from Illumina for use with the Nextera XT kit. Final indexed products (0.8x bead:pool ratio) were pooled to an equimolar concentration of 0.5-0.8nM. The Agilent High Sensitivity DNA Kit for Bioanalyzer 2100 (Agilent Technologies) was used to confirm amplicon presence, size and adapter-dimer removal. The cleaned pool was sequenced on an Illumina MiSeq sequencing platform with a 600- cycle V3 kit. All sequencing was performed at the Teagasc Sequencing Facility, per standard Illumina protocols.

### Optimised RNA extraction to semi-nested PCR method

In order to improve the RNA yield from naturally contaminated shellfish, variations with regard to sample extract volume were included. RNA was extracted from 1000 μl of proteinase K extract and eluted into a smaller volume (30 μl). All samples were extracted in duplicate. LunaScript and AmpliTaq Gold provided the highest-quality HTS reads with minimum errors. RT with LunaScript was performed in triplicate to provide sufficient cDNA for semi-nested PCR of norovirus GI and GII targets in triplicate. cDNA was pooled and stored at -20°C. The first round of semi-nested PCR was prepared as follows, 10 µl cDNA per technical triplicate was added to a 45 μl reaction mixture with a final concentration of 10 mM Tris-HCl (pH 8.3), 50 mM KCl, 20 μM of dNTPs, 2 μM of each primer (see Table 10), 2.5mM of MgCl2 and 2.5 U of AmpliTaq® DNA Polymerase (Applied Biosystems, USA). The first round PCR product (5 µl) was subsequently added to 45 µl of a reaction mixture containing 10 mM TrisHCl (pH 8.3), 50 mM KCl, 20 μM of dNTPs, 0.4 μM of each primer, 2.5 mM of MgCl2 and 2.5 U of AmpliTaq® DNA Polymerase. The Agilent, High Sensitivity DNA Kit for Bioanalyzer 2100 (Agilent Technologies), was used to visualise second-round PCR products. Following library preparation, an additional Ampure bead clean-up step (0.7x bead:pool ratio) was performed to remove adapter dimers and 1 μl of the cleaned pool was visualised using the Agilent High Sensitivity DNA Kit for Bioanalyzer 2100 (Agilent Technologies) to confirm adapter-dimer removal.

### Bioinformatic analysis

The pipeline utilised is based on the results from a previous study, which benchmarked pipelines and classifiers for norovirus amplicon analysis (81). Adapters and primers were trimmed using cutadapt (v 2.6) with an -e 0.1 and a minimum length of 100 bp. Reads were quality filtered in VSEARCH (v2.4.2) with a minimum length of 100 bp and a maximum length of 400 bp, a minimum overlap of 50 bp, a maximum of 20% mismatches in the alignment and a maximum expected error threshold of 1. Chimera removal was performed using UCHIME within VSEARCH (v2.4.2) using *de novo* and reference-based chimera removal, with 99% clustering prior to chimera detection. The database for chimera-based removal was generated as follows; all available norovirus sequences greater than 1000 bp were fetched from GenBank using rentrez (v1.2.3). VP1 sequences were created using the second-round primers outlined in Table 10 in seqkit (v1.4), and sequences were clustered to 85% identity using CD-HIT (v4.7). Clustering of the sequences following chimera removal was performed at 99% identity, with a minimum of 1 read per sample required for a true sequence. OTUs representing less than 1% of reads per sample were removed. OTUs were classified using the NoroNet typing tool from RIVM.

### Statistical analysis

All analysis was performed using R statistics (v 4.2.1) in R Studio. Kruskal-Wallis tests were performed in base R, while the post hoc test for Kruskal-Wallis was conducted using the Dunn test in the R package rstatix (v 0.7.0) (82). Factor Analysis of Mixed Data (FAMD) was performed using R package factoextra (v 1.7.0) (83) and factoMineR (v2.6.0) (84). DNA polymerase and RT enzyme were included as the factors of interest, alongside the numeric variable of interest such as mean Phred score or mean expected errors. Kendall correlation tests were performed using R statistics and interpreted based on previously reported ranks (34).

Distance matrices were conducted using the R package vegan (v 2.6.2). All distance measures were conducted using 999 permutations (Jaccard). Analysis of variance using distance matrices (ADONIS2) was also performed using the vegan package in R with Bonferroni p-value correction. Post Hoc tests for ANOSIM/ADONIS2 were performed using the R package RVAideMemoir (v 0.9-81-2).

A confusion matrix was generated using the yardstick package (v 1.0.9) in R from tidymodels (85). Data was coded in a binary fashion, encoding 1 for agreement between expected and observed data and zero for disagreement. Classifiers and databases were compared based on the sensitivity or true positive rate (TPR), false positive rate (FPR) or 1-specificity, F1 score and balanced accuracy (average of sensitivity and specificity). Sensitivity refers to the probability of obtaining a positive test for a true positive, and false positivity rate refers to the probability of obtaining a false positive test for a true negative or, in this case misclassification or missed classification. F1 score takes the harmonic mean of the sensitivity and specificity, while balanced accuracy takes the mean of the sensitivity and specificity. Jaccard distance measures (R package vegan v 2.6.4) were used to assess true and false matches, between expected and observed data.

For UniFrac analysis, files were imported into QIIME2/2021.2. Sequences were aligned using the MAFFT plugin and masked. For UniFrac analysis, rooted trees were generated using rooted fastree and distances computed with all tips.

Custom BLAST databases were created based on the expected data for each library 1 and 2. Observed output for each sample was blasted against the custom database, requiring 99% similarity at 75% coverage of the amplicon. Multiple hits for an observed sequence to a reference sequence in the BLAST DB were filtered. The observed OTU with the highest bit-score and lowest e-value per sample/library was selected for the comparison if multiple hits were obtained.

### Data availability

Sequence data generated during the current study have been deposited in the European Nucleotide Archive under accession number PRJEB58629.

## Data Availability

The scripts used for the processing of bioinformatic data are available at https://github.com/ahfitzpa/Norovirus_HTS_amplicon. The dataset generated and analysed during the current study is available in the ENA repository under accession number PRJEB58629.

## Author Contributions

AHF, AR and SK designed the experiments. AR prepared the spiked samples, from RNA extraction to sample characterisation using Sanger sequencing. AHF performed RT-PCRs, library preparation and bioinformatic analysis and wrote the manuscript. FC sequenced all libraries at the Teagasc Next Generation DNA Sequencing Facility. SK, HO’S and PC reviewed the final draft. All authors contributed to the article and approved the submitted version.

## Acknowledgements

Figures 1–2, and Supplementary Figure 1 were created with BioRender.com. The authors would like to thank Leon Devilly and James Fahy (Shellfish Microbiology team, Marine Institute) for their assistance with sample preparation for sequencing, Elaine Lawton and Teagasc Next Generation DNA Sequencing Facility for supporting this work.

---

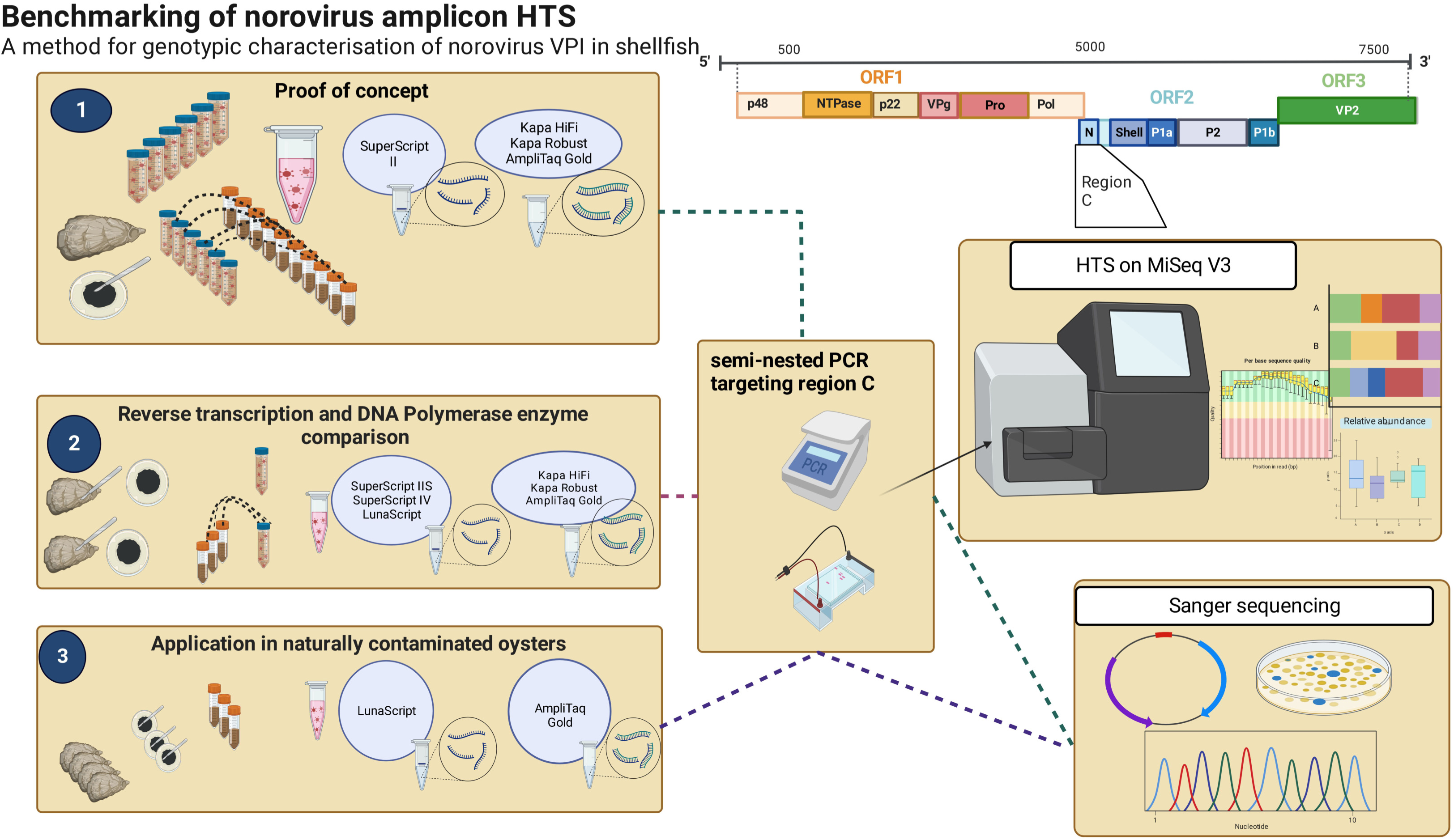

